# Increased neuronal activity restores circadian function in *Drosophila* models of C9orf72-ALS/FTD

**DOI:** 10.1101/2025.08.07.669085

**Authors:** Sho Inami, Benjamin P. Jenny, Oghenerukevwe Akpoghiran, Sara I. Gallagher, Alexandra K. Strich, Ayako Tonoki, Davide Trotti, Aaron R. Haeusler, Kyunghee Koh

## Abstract

Circadian rhythm disruptions are common across neurodegenerative diseases, but their link to amyotrophic lateral sclerosis (ALS) and frontotemporal dementia (FTD) remains unclear. The *C9orf72* hexanucleotide repeat expansion is the most prevalent genetic cause of ALS/FTD. Here, we used *Drosophila* models expressing toxic arginine-rich dipeptides (PR or GR) or GGGGCC hexanucleotide repeats to investigate circadian deficits in C9orf72-ALS/FTD. We found that circadian rhythmicity and period length were disrupted in a repeat number-, dosage-, and age-dependent manner. Additionally, we observed lower levels of the neuropeptide PDF, a key regulator of free-running circadian rhythms, as well as decreased projection complexity and reduced neuronal activity in PDF-expressing neurons. Importantly, increases in neuronal activity significantly restored circadian function under select conditions. These results implicate reduced neuronal activity in C9orf72-ALS/FTD circadian deficits, underscoring the importance of precisely tuned, circuit- and stage-specific interventions.

**Highlights:**

- C9orf72 dipeptide and nucleotide repeats disrupt circadian rhythms in *Drosophila*
- Circadian dysfunction with reduced PDF and neurites emerges before neuron loss
- Increased neuronal activity rescues mild circadian dysfunction
- Activity-based rescue is effective across ages and models when precisely tuned

## Introduction

Growing evidence links circadian rhythm disruptions to neurodegenerative disorders, including Alzheimer’s and Parkinson’s diseases, where circadian dysfunction often precedes and potentially exacerbates neurodegeneration^1–4^. In contrast, the relationship between circadian disturbances and amyotrophic lateral sclerosis (ALS) or frontotemporal dementia (FTD) remains less well defined. Some evidence supports such a link: ALS mouse models harboring mutations in *SOD1* show altered expression of core clock genes^5^, and structural MRI studies reveal hypothalamic atrophy, a region encompassing the circadian pacemaker and sleep-regulatory centers, in both ALS patients and presymptomatic gene carriers^6^. Moreover, a recent study found altered sleep architecture, including delayed sleep onset, in presymptomatic carriers of familial ALS/FTD mutations and in early-stage patients^7^. The delayed sleep onset could result from reduced sleep drive but may also reflect disrupted circadian timing. Thus, although prior studies suggest a link between ALS/FTD and circadian dysfunction, experimental models directly investigating this relationship have been lacking, which has limited our understanding of how ALS/FTD-related pathology impacts circadian rhythms.

ALS is characterized by progressive degeneration of motor neurons, while FTD involves degeneration of cortical neurons, leading to cognitive and behavioral impairments. These two conditions form a clinical and pathological continuum with overlapping genetic causes. The most common genetic contributor to familial ALS and FTD is a hexanucleotide (GGGGCC, or G_4_C_2_) repeat expansion (HRE) in the *C9orf72* gene, accounting for ∼40% of familial and 5–10% of sporadic ALS cases^8–10^. This *C9orf72* HRE contributes to disease via multiple mechanisms: sequestration of RNA-binding proteins by RNA foci, repeat-associated non-AUG (RAN) translation of toxic dipeptide repeat proteins (DPRs), especially the arginine-rich PR and GR species, and reduced C9orf72 protein levels^11–18^. *Drosophila melanogaster*, despite lacking a *C9orf72* homolog, offers a genetically tractable, *in vivo* system to model the toxic gain-of-function mechanisms of DPRs and G_4_C_2_ repeats.

An important but incompletely understood mechanism in ALS/FTD pathogenesis is altered neuronal activity. Studies in C9orf72-ALS/FTD patient-derived neurons and mouse models have reported both increased and decreased excitability^19^. Although the origins of these mixed findings are not well understood, several factors such as neuronal age, culture duration, and the presence of glia, may influence the excitability phenotypes (Sareen et al., 2013; Wainger et al., 2014). Some patient-derived neuron studies suggest a biphasic trajectory, with early hyperexcitability transitioning to hypoexcitability at later stages^20^. This pattern resembles findings in Alzheimer’s disease patients, where cortical and hippocampal hyperactivity in preclinical stages shifts toward hypoactivity as neurodegeneration progresses^23–26^. To better understand the functional impact of excitability changes *in vivo*, we examined whether manipulating neuronal activity could modulate circadian behavior in *Drosophila* models of *C9orf72*-linked ALS/FTD.

Circadian rhythms are driven by molecular clocks consisting of transcriptional-translational feedback loops that generate ∼24-hour oscillations in gene expression^27,28^. In mammals, the master circadian clock resides in the suprachiasmatic nucleus (SCN) of the hypothalamus. In *Drosophila*, rhythmic behavior is governed by a network of ∼150 clock neurons, among which the small ventral lateral neurons (s-LNvs), which express the neuropeptide pigment-dispersing factor (PDF), are crucial for maintaining free-running circadian rhythms in constant darkness^29–31^.

Here, we used the binary Gal4/UAS system to express toxic PR, GR, or G_4_C_2_ repeats selectively in PDF+ s-LNvs and assessed their effects on circadian behavior. We found that these constructs induced distinct circadian phenotypes depending on repeat type, length, dosage, and age. Behavioral disruptions occurred before detectable cell loss and were accompanied by reduced PDF expression and simplified dorsal projections. Our findings reveal that increases in neuronal activity can restore circadian function in multiple C9orf72 models under conditions of relatively mild dysfunction. We did not observe any condition in which enhancing neuronal activity exacerbated circadian phenotypes, suggesting that a biphasic excitability profile may not apply to circadian neurons. Together, our findings indicate that reduced neuronal activity contributes to circadian deficits in C9orf72-ALS/FTD and suggest that therapeutic strategies targeting excitability may require careful tuning to cell type and disease stage.

## Results

### Expression of C9orf72 DPRs and nucleotide repeats impairs circadian rhythmicity and alters period length

To assess the impact of C9orf72 DPRs and G_4_C_2_ repeats on *Drosophila* circadian behavior, we used the Gal4/UAS binary expression system to express PR, GR, or G_4_C_2_ repeats in PDF-positive (PDF+) clock neurons (Figure 1A). The UAS-(PR)_36_ construct encodes 36 copies of the PR dipeptide, while the UAS-(GR)_100_ construct expresses 100 copies of the GR dipeptide^32^. The UAS-(G_4_C_2_)_49_ construct generates 49 G_4_C_2_ repeats, potentially producing GR, GA, and GP dipeptides from the sense strand, as well as forming RNA foci^33^. Expression of any of these constructs in circadian pacemaker neurons using the *Pdf*-Gal4 driver significantly reduced circadian rhythm strength in both male and female flies (Figure 1B). Notably, (PR)_36_ expression led to a significant lengthening of the circadian period compared to controls, whereas (GR)_100_ expression caused a modest but significant shortening of period length (Figure 1B). (G_4_C_2_)_49_ expression also slightly reduced the circadian period, although this effect was not statistically significant. These findings suggest that C9orf72-related constructs differentially disrupt core circadian function, with phenotypic outcomes influenced by repeat type and length. For the remainder of the study, we focused on male flies, which exhibited more robust rhythmicity than females in our control genotypes. This sex difference is consistent with previous reports showing reduced rhythmicity in mated females compared to males in the iso31 genetic background used in this study^34^.

**Figure 1:**
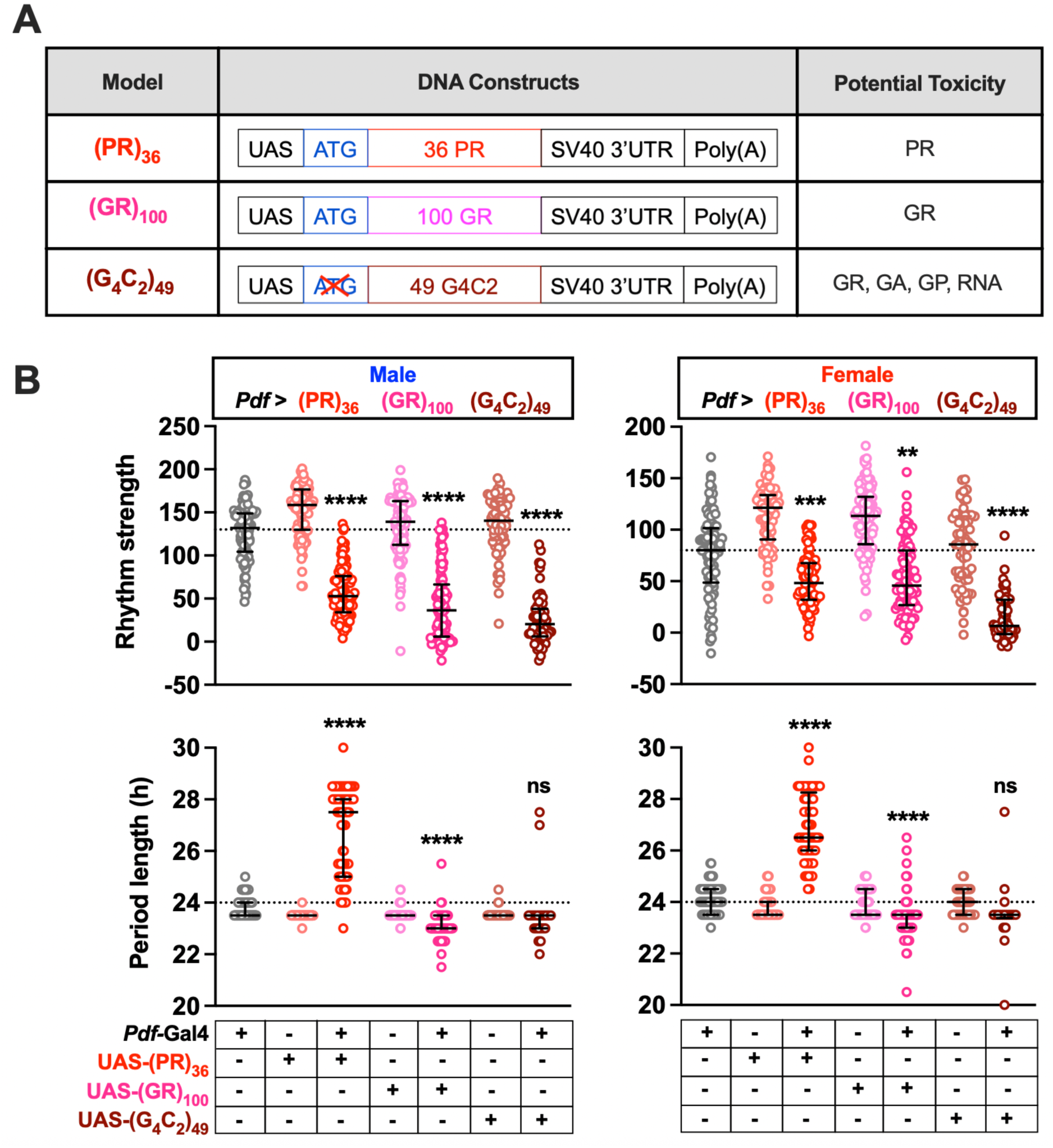
Expression of (PR)_36_, (GR)_100_, and (G_4_C_2_)_49_ in PDF+ clock neurons results in reduced rhythm strength and altered period length. (A) Schematic diagram of the UAS constructs used and their potential toxicity. (B) The rhythm strength (top) and period length (bottom) of mated males (left) and females (right) of indicated genotypes. The median and interquartile range are shown. N = 64–111 (male) and n = 62-94 (female) for rhythm strength; n = 18–88 (male) and n = 24–87 (female) for period length. Ns are lower for period length because only rhythmic flies are included. The horizontal dotted line represents the median rhythm strength and period length of the common control (*Pdf*-Gal4*/+*). Statistical comparisons shown represent the less significant of the two pairwise tests (each genotype expressing a C9orf72 construct was compared to both parental controls: *Pdf*-Gal4/+ and UAS-transgene/+). **p<0.01, ***p < 0.001, ****p < 0.0001, ns: not significant; Kruskal–Wallis test followed by Dunn’s multiple comparisons test relative to parental controls.

### Repeat length, dosage, and age influence the severity of circadian disruption

A notable observation from our dataset is that (GR)_100_ or (G_4_C_2_)_49_ expression in PDF+ neurons exhibited significantly reduced rhythmicity and slightly shortened period length, circadian phenotypes resembling those seen in *Pdf*-null mutants or flies in which PDF+ neurons have been ablated^31^. These findings suggest that (GR)_100_ and (G_4_C_2_)_49_ elicit particularly severe circadian disruption, likely representing the maximal phenotype achievable through perturbation of PDF+ neurons. In contrast, the reduced rhythm strength and lengthened period seen with (PR)_36_ expression may reflect a milder degree of circadian dysfunction.

To determine whether the distinct phenotypes reflect differences in repeat length, dosage, or type, we first compared PR constructs of different lengths. Compared to (PR)_36_, (PR)_100_ expression led to a more severe reduction in rhythm strength and a shift to shorter periods (Figure 2A), resembling the effects of (GR)_100_ and (G_4_C_2_)_49_ expression. We next tested dose dependence by comparing one versus two copies of UAS-(PR)_36_. Doubling the (PR)_36_ dosage further decreased rhythm strength and resulted in a shortened period, again phenocopying (GR)_100_ and (G_4_C_2_)_49_ (Figure 2B). These results suggest that the long-period phenotype observed with one copy of (PR)_36_ represents a relatively mild disruption of the circadian system, and increasing the repeat length or dosage pushes the system toward more severe disruption similar to that induced by (GR)_100_ and (G_4_C_2_)_49_ toxicity.

**Figure 2:**
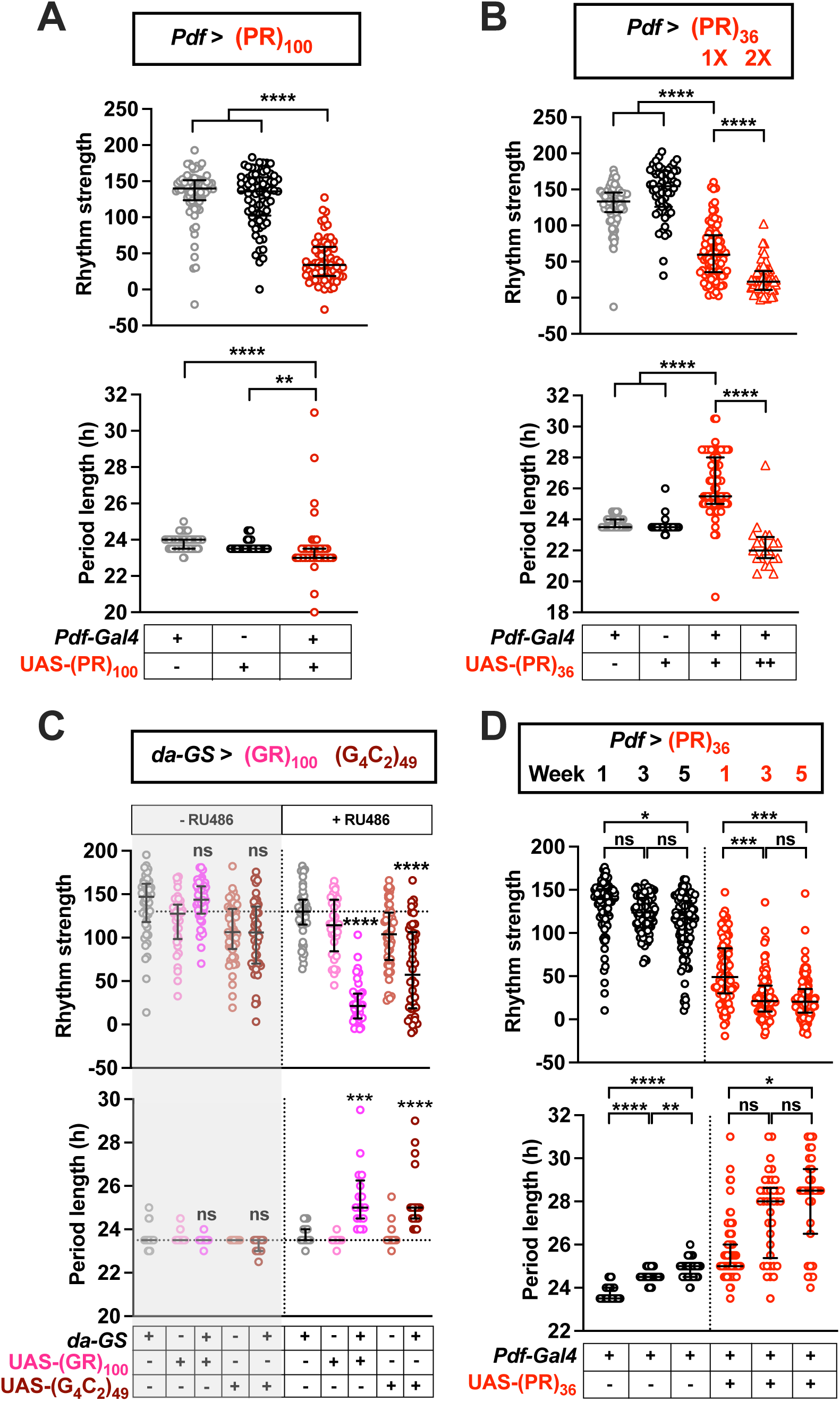
Repeat length, dosage, and age influence the severity of C9orf72-linked circadian disruption. **(A)** Rhythm strength and period length of males expressing (PR)_100_ in PDF+ neurons compared to parental controls. N = 69–78 for rhythm strength; n = 39–77 for period length. **(B)** Rhythm strength and period length of males expressing one or two copies of (PR)_36_ and controls. N = 56–95 for rhythm strength; n = 20–94 for period length. **(C)** Rhythm strength and period length of males expressing (GR)_100_ or (G_4_C_2_)_49_ ubiquitously in the adult (using *da*-GS) and their parental controls. Flies were fed 50 μM RU486 or vehicle (1% ethanol). The horizontal dotted line represents the median rhythm strength and period length of the common control (*da*-GS */+*) for the +RU486 condition. N = 35–48 for rhythm strength; n = 17–48 for period length. **(D)** Rhythm strength and period length of control (*Pdf > +*) and (PR)_36_-expressing (*Pdf >* (PR)_36_) male flies. For the 1-week condition, circadian assays were performed on flies aged 3–4 days at the start of the assay. For the 3-week and 5-week conditions, the flies were 3 and 5 weeks old, respectively, at the start of the assay. N = 92–107 for rhythm strength and n = 31–99 for period length. For all panels, data are shown as median and interquartile range. *p < 0.05, **p < 0.01, ***p < 0.001, ****p < 0.0001, ns: not significant; Kruskal–Wallis test followed by Dunn’s multiple comparisons test of indicated pairwise comparisons (A, B, D) or relative to both parental controls (C).

To determine whether the period lengthening is specific to PR toxicity or a general feature of lower toxicity levels, we used the GeneSwitch (GS) drug-inducible system, which permits temporal and dosage control of transgene expression, to mildly express (GR)_100_ and (G_4_C_2_)_49_ in adult flies. Unlike the *Pdf*-Gal4 driver, which is active in PDF+ clock neurons beginning in the third-instar larval stage^35^, *daughterless*-GeneSwitch (*da*-GS) drives ubiquitous expression in response to RU486 feeding. We activated the driver by feeding adult flies 50 µM RU486 starting at 3–4 days of age. Under these conditions, both (GR)_100_ and (G_4_C_2_)_49_ expression led to decreased rhythm strength (Figure 2C). Importantly, both also caused an increase in period length, recapitulating the phenotype observed with (PR)_36_ (Figures 1 and 2B). These findings demonstrate that period lengthening is not exclusive to PR expression, but may reflect mild circadian dysfunction, regardless of repeat type.

### Aging exacerbates (PR)_36_-induced circadian dysfunction

Given that aging is a major risk factor for neurodegenerative diseases^36,37^, we next investigated how the mild (PR)_36_-induced circadian phenotypes change with age. We assessed rhythm strength and period length of flies expressing (PR)_36_ or control flies at three ages: “one week”, three weeks, and five weeks, corresponding to 3–4 days, 21–22 days, or 35–36 days old at assay start. In control flies, rhythm strength declined gradually with age. Five-week-old controls showed a significant reduction in rhythm strength relative to one-week-old flies, while the decrease at three weeks did not reach significance (Figure 2D). In contrast, (PR)_36_-expressing flies exhibited a more rapid decline in rhythm strength, with significant reductions already apparent at three weeks, with no further decrease at five weeks, consistent with a floor effect (Figure 2D). These findings indicate that while aging impairs rhythm strength in both control and (PR)_36_ flies, the decline occurs earlier in (PR)_36_-expressing flies.

We also observed age-related changes in period length. In control flies, period length increased significantly with age, consistent with previous findings^38^. A similar pattern was observed in (PR)_36_-expressing flies; period length was significantly longer in five-week-old flies compared to one-week-old counterparts (Figure 2D). Notably, the age-dependent period lengthening differs from the period shortening observed with increased PR dosage, longer repeats, or expression of (GR)_100_ and (G_4_C_2_)_49_. This difference is unlikely to reflect weaker effects of aging, as rhythm strength in five-week-old (PR)_36_ flies was as low as that seen with the more toxic C9orf72 constructs. Instead, aging may exert a stronger influence on rhythm strength than on period length, a possibility we consider further in the Discussion.

### Disrupted circadian behavior precedes cell loss in C9orf72 models

To determine whether the circadian behavioral phenotypes caused by (PR)_36_, (GR)_100_, and (G_4_C_2_)_49_ expression are due to the loss of PDF+ neurons, we assessed the survival of these clock cells in adult fly brains. We examined flies ∼10 days post-eclosion, corresponding to the endpoint of the 1-week behavioral circadian assays. To visualize PDF+ neuron survival, we expressed the nuclear marker RedStinger^39^ under the control of *Pdf*-Gal4, which labels 8 PDF+ neurons per hemisphere: 4 small ventrolateral neurons (s-LNvs) and 4 large ones (l-LNvs). We focused our analysis on the s-LNvs, which are essential for maintaining circadian rhythms in constant darkness, in contrast to the l-LNvs, which are dispensable for free-running behavior ^31^. At this early time point, there were no significant differences in the number of s-LNvs across genotypes, (Figures 3A and 3B, top row), including in flies expressing (GR)_100_ or (G_4_C_2_)_49_ (p > .33). Since the severity of rhythm deficits induced by (GR)_100_ and (G_4_C_2_)_49_ at this time point is comparable to that of flies with ablated s-LNvs, these findings suggest that pronounced behavioral effects can arise in the absence of neuron loss.

**Figure 3:**
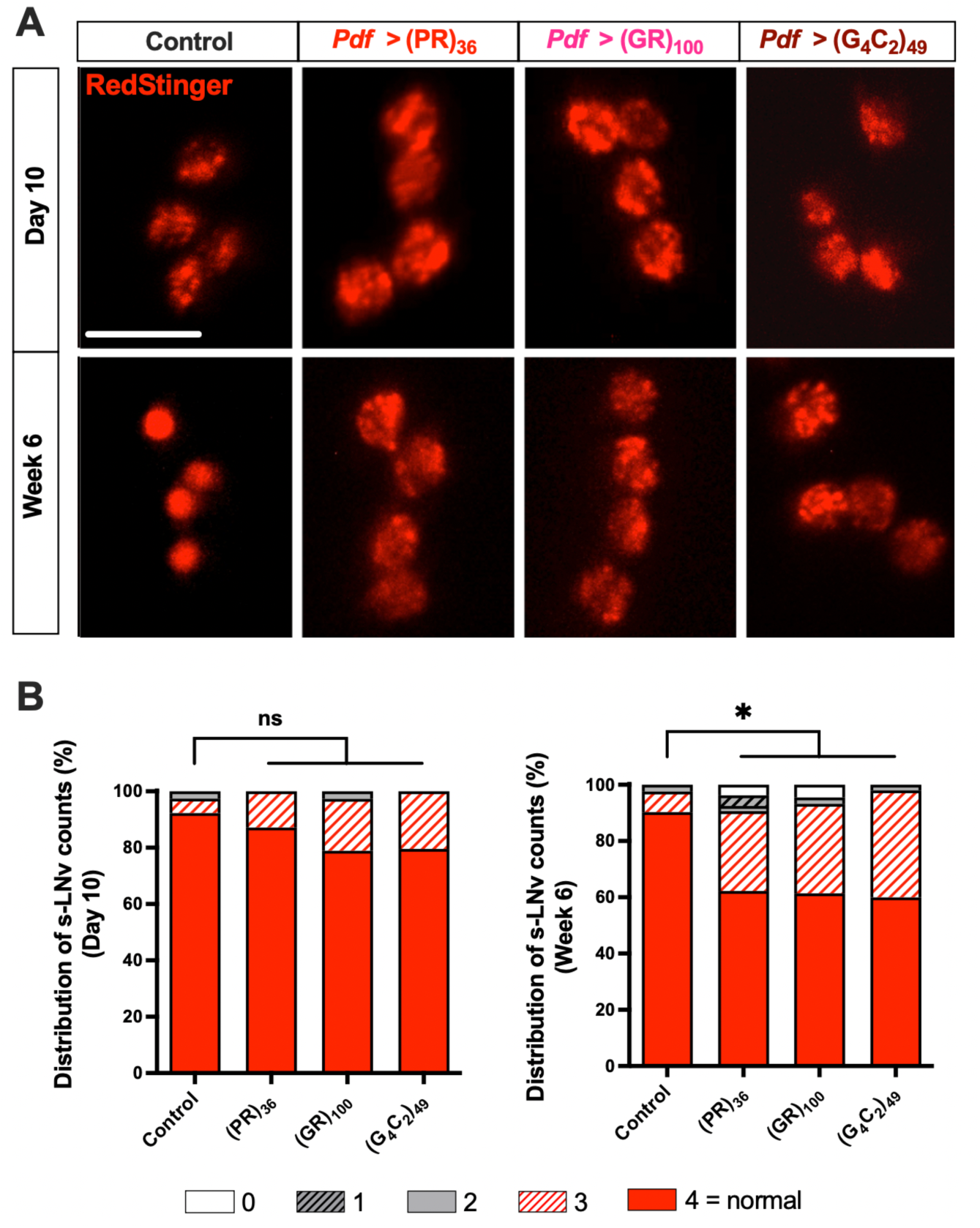
Circadian dysfunction precedes neuronal loss in C9orf72 models. **(A)** Representative expression pattern of the RedStinger nuclear marker in the PDF+ s-LN_v_s of control flies and flies expressing (PR)_36_, (GR)_100_, (G_4_C_2_)_49_ at ∼10 days (top) or ∼6 weeks (bottom) of age. Modest but significant cell loss is detected at 6 weeks, but not at 10 days. Scale bar: 10 μm. **(B)** Number of RedStinger-expressing s-LNvs in each hemibrain. Percentage of hemibrains with indicated number of s-LNvs is shown. N = 38–49 (Day 10); n = 41–53 (Week 6). *p < 0.05, ns: not significant; Kruskal-Wallis test followed by Dunn’s multiple comparisons test.

Since aging exacerbated the behavioral phenotypes in (PR)_36_-expressing flies (Figure 2D), we next asked whether neuronal loss occurs later. We quantified the number of s-LNvs at ∼6 weeks post-eclosion, the endpoint of the 5-week circadian assays. Despite the overall viability of most PDF+ neurons (Figures 3A, bottom row), all three C9orf72 constructs caused a significant reduction in s-LNv number compared to controls by this age (Figure 3B, bottom row). Together, these results demonstrate that circadian dysfunction in C9orf72-expressing flies emerges before substantial neuron loss, and likely reflects early deficits in neuropeptide signaling and structural integrity rather than cell death per se.

### Decreased PDF expression and neuronal processes in the s-LNvs in C9orf72 models

Flies lacking PDF exhibit arrhythmic behavior and slightly shortened circadian periods^31^, resembling the effects of (GR)_100_ and (G_4_C_2_)_49_ expression. We therefore investigated whether these disruptions might result from reduced PDF expression in the s-LNvs. In wild-type flies, PDF levels cycle daily in the dorsal projections of the s-LNvs^40,41^. To assess whether PDF expression is altered, we performed immunostaining under light–dark (LD) conditions at Zeitgeber Time 2 (ZT2), a time point when PDF levels in the dorsal projections peak and are minimally influenced by differences in circadian period length. PDF levels in the s-LNv cell bodies were significantly reduced across all three C9orf72-expressing groups (Figures 4A, top row, and 4B). In the dorsal projections, (GR)_100_ and (G_4_C_2_)_49_-expressing flies showed marked reductions in PDF immunoreactivity, while (PR)36 flies exhibited a milder, non-significant decrease (Figures 4A, middle row, and 4C).

**Figure 4:**
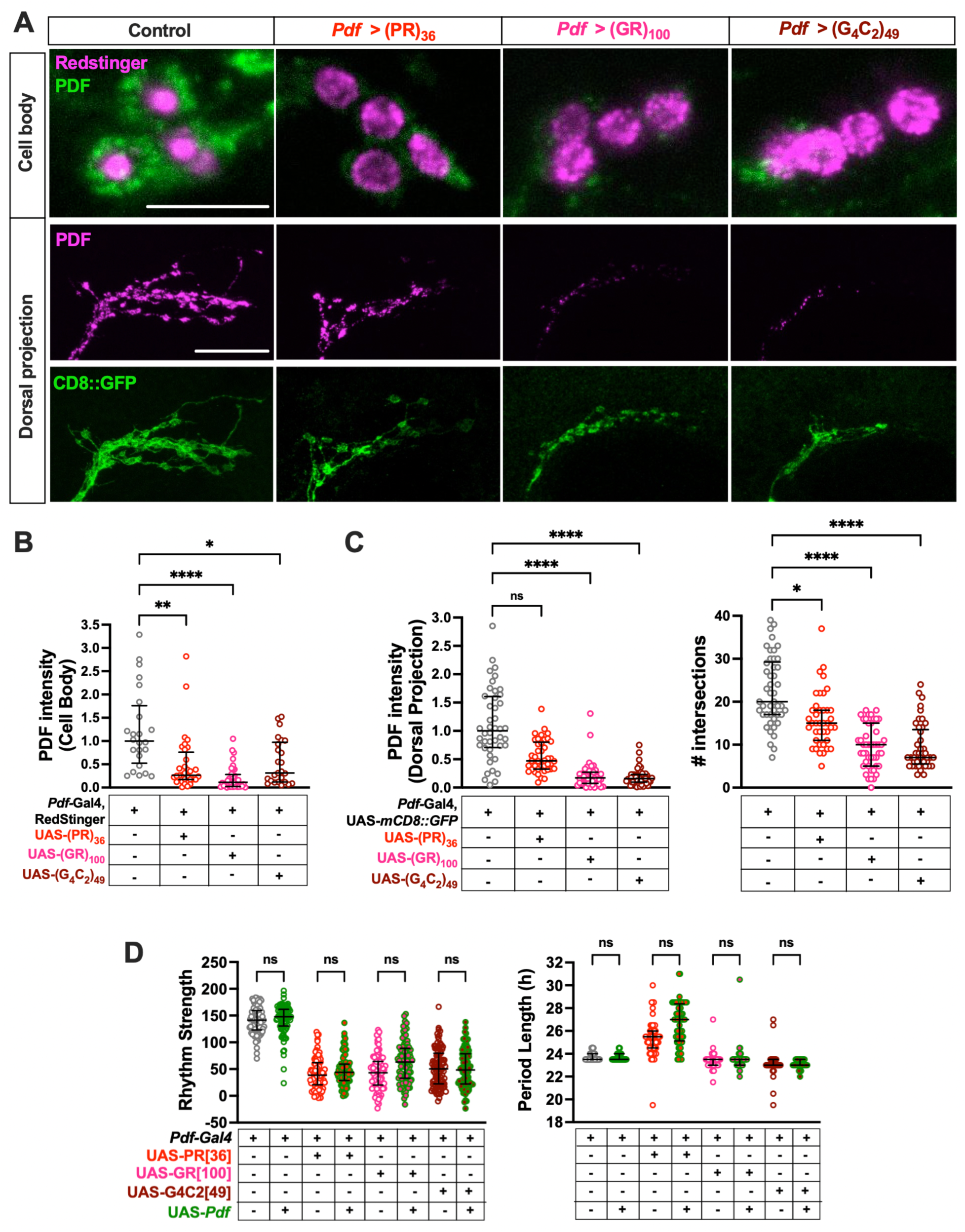
Expression of C9orf72 transgenes reduces PDF levels and projection complexity in s-LNv neurons, and PDF overexpression does not restore rhythmicity. **(A)** Representative confocal images showing PDF expression in s-LNv cell bodies (top) and dorsal projections (middle), and CD8::GFP-labeled dorsal projections (bottom) in control flies and flies expressing (PR)_36_, (GR)_100_, or (G_4_C_2_)_49_. RedStinger was used to identify s-LNv cell bodies, as PDF expression was frequently undetectable in this subset of neurons in C9orf72-expressing brains. CD8::GFP images of C9orf72-expressing brains were brightened in Fiji to facilitate quantification of projection complexity. Scale bar: 10 μm (cell bodies); 20 μm (dorsal projections). **(B)** Quantification of PDF levels in s-LNv cell bodies. N = 24–36 for cell bodies; n = 41-47 for dorsal projections. **(C)** Quantification of PDF levels in s-LNv dorsal projections (left) and Scholl analysis of s-LNv dorsal projections (right), in which the number of intersections between the projections and concentric circles spaced 10 μm apart was quantified. N = 40–47. **(D)** Rhythm strength and period length of control males (*Pdf*-Gal4/+) and males expressing indicated C9orf72 constructs with or without co-expression of PDF. N = 86–95 for rhythm strength; n = 61–91 for period length. For all plots, data are presented as median and interquartile range. *p < 0.05, **p < 0.01, ****p < 0.0001, ns: not significant; Kruskal-Wallis test followed by Dunn’s multiple comparisons test.

To assess whether the reductions in PDF in the s-LNv dorsal projections were accompanied by structural changes, we co-stained with membrane-targeted CD8::GFP to visualize neuronal morphology, which normally exhibits daily remodeling and exhibits peak complexity and openness in the early morning^40^. Since expression of all three C9orf72 constructs significantly reduced CD8::GFP signal intensity in the dorsal projections compared to controls, we adjusted image brightness in Fiji to facilitate quantification of the dorsal projection complexity. Visual inspection suggested that s-LNv dorsal projections were less complex in C9orf72-expressing flies (Figure 4A, bottom row). We quantified these morphological changes using Scholl analysis, measuring the number of intersections between dorsal processes and concentric rings spaced 10 μm apart. (GR)_100_ and (G_4_C_2_)_49_ expression led to a significant reduction in intersection counts, while (PR)_36_ produced a milder effect (Figure 4D). As no s-LNv neuron loss was observed at this stage, the reduction in PDF and projection complexity likely reflects the onset of neurodegenerative processes prior to overt cell death.

Given the reduction in PDF levels in C9orf72-expressing flies, we asked whether increasing PDF expression could restore circadian function. To test this, we used a UAS-*Pdf* transgene to overexpress PDF in the LNvs. We found that this manipulation did not significantly improve the circadian phenotypes in any of our C9orf72 models (Figure 4D). These results suggest that although reduced PDF levels likely contribute to circadian dysfunction, PDF overexpression alone is insufficient to restore normal circadian function. Together, these findings suggest that C9orf72-associated toxicity disrupts both neuropeptide expression and the structural integrity of core pacemaker neurons, and that restoring PDF alone is insufficient to rescue circadian function, highlighting the complexity of the mechanisms underlying neuronal dysfunction in these models.

### Neural activity in PDF+ neurons modulates the circadian phenotypes of (PR)_36_ expression

Altered neuronal excitability is a frequently reported feature of ALS pathophysiology, observed in both patient-derived neurons and animal models^20–22,42,43^. To increase excitability, we co-expressed TrpA1, a warmth-sensitive cation channel that becomes active at temperatures above 25°C and increases neuronal firing^44^. In a pilot experiment, TrpA1 activation at 27°C resulted in severely reduced circadian rhythmicity in control flies without altering the phenotype of (PR)_36_-expressing flies. To avoid confounding effects of strong TrpA1 activation, we reared and tested flies at 25°C, where the channel is only weakly active. At this temperature, TrpA1 activation alone led to a modest decrease in rhythm strength, but its co-expression with (PR)_36_ significantly rescued the circadian phenotype; rhythm strength improved, and period length shortened compared to (PR)_36_ expression alone (Figure 5A).

**Figure 5:**
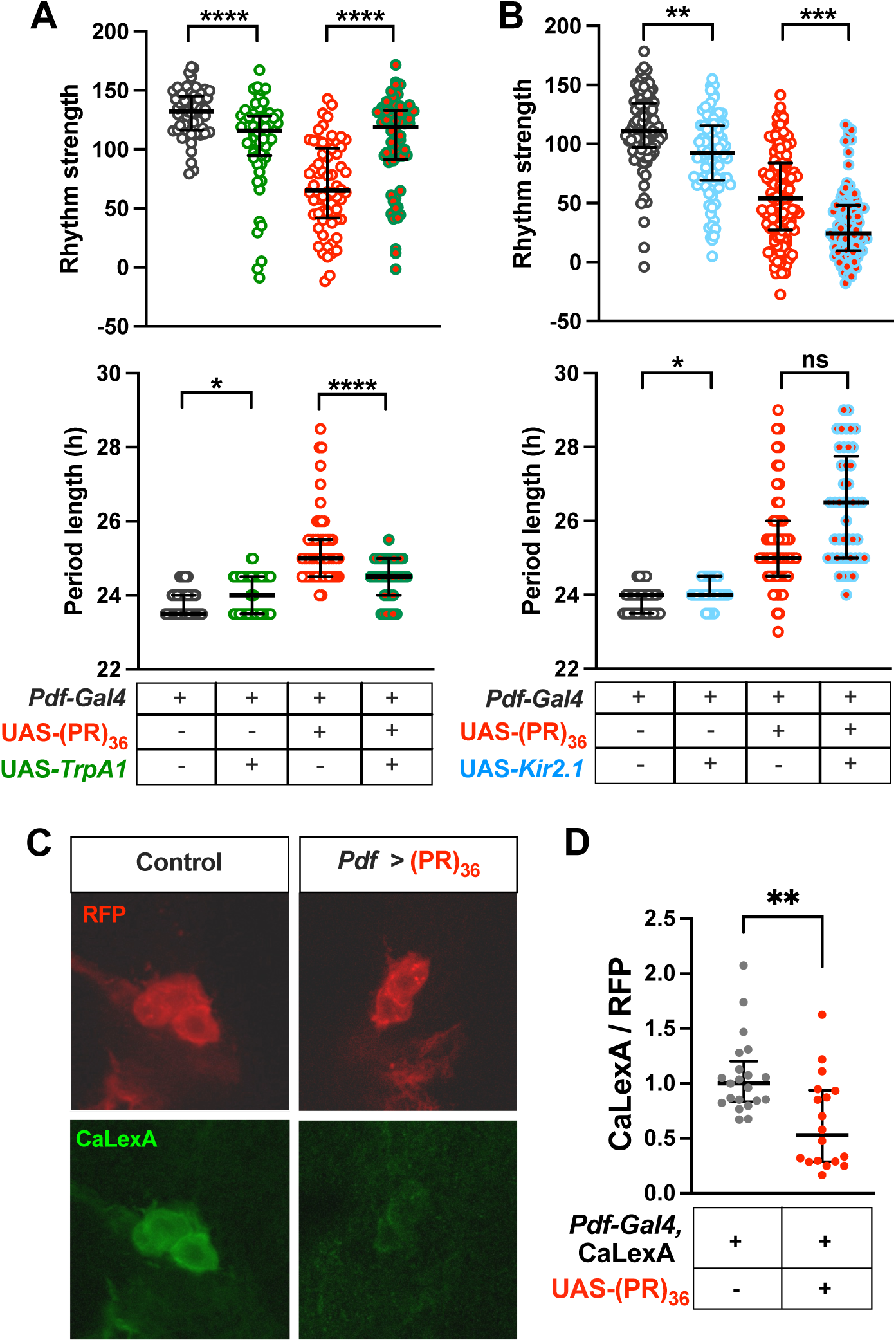
Neuronal activity modulates (PR)_36_-induced circadian phenotypes. (**A)** Rhythm strength (top) and period length (bottom) of males of the indicated genotypes. N = 55–64 for rhythm strength; n = 55–57 for period length. **(B)** Rhythm strength (top) and period length (bottom) of males of the indicated genotypes. N = 88–128 for rhythm strength; n = 45–99 for period length. **(C)** Representative confocal images showing RFP (internal control) and CaLexA (calcium reporter) expression in the s-LNvs of control flies and flies expressing (PR)_36_. **(D)** Normalized CaLexA signal relative to RFP internal control. N = 18-21. For all plots, data are presented as median and interquartile range. *p < 0.05, **p < 0.01, ***p < 0.001, ****p < 0.0001, ns: not significant; Kruskal-Wallis test followed by Dunn’s multiple comparisons test.

Next, to test the effects of decreasing excitability, we used Kir2.1, an inward-rectifying potassium channel that hyperpolarizes neurons and suppresses firing^45^. Kir2.1 expression alone slightly reduced rhythm strength. However, when combined with (PR)_36_, Kir2.1 further exacerbated circadian dysfunction, leading to a pronounced reduction in rhythm strength and a significant lengthening of period (Figure 5B). These results indicate that neuronal excitability modulates (PR)_36_-induced circadian disruption; increasing excitability restores circadian function, while decreasing it worsens dysfunction.

To test whether (PR)_36_ expression leads to decreased neural activity, we used CaLexA, a GFP-based reporter of calcium-dependent neural activity^46^, in adult flies aged ∼10 days, corresponding to the age at the end of our 1-week behavioral assay. Co-expressed RFP served as an internal control. We found that (PR)_36_-expressing flies significantly reduced CaLexA signal in PDF+ neurons, suggesting decreased calcium-dependent neural activity (Figures 5C and 5D). These results support the idea that (PR)_36_ expression reduces neuronal excitability in circadian pacemaker neurons, contributing to the observed behavioral deficits.

### Increased neuronal activity can restore circadian function at different ages and across multiple C9orf72 models

Given that aging exacerbates (PR)_36_-induced circadian dysfunction, we next asked whether the beneficial effects of neuronal activation via TrpA1, observed in young flies, could also rescue circadian defects in older animals. To assess this, we conducted circadian assays in flies aged 1 and 3 weeks and evaluated the efficacy of excitability-based intervention. Under chronic activation conditions (Figure 6A), flies were reared, aged, and tested at 25°C, a temperature that induces low-level TrpA1 activity, as in our earlier rescue experiment in young flies. In this setting, expression of (PR)_36_ and TrpA1 under *Pdf*-Gal4 began at the third-instar larval stage^35^. As shown above (Figure 2D), both control and (PR)_36_-expressing flies exhibited age-dependent declines in rhythm strength and increases in period length (Figure 6B). A similar decline in rhythm strength was observed in flies expressing TrpA1 alone, suggesting that prolonged activation may exert cumulative negative effects on clock neurons. At one week of age, co-expression of TrpA1 with (PR)_36_ restored rhythm strength and normalized period length, replicating our finding shown in Figure 5A. However, this rescue effect was lost in 3-week-old flies; TrpA1 co-expression no longer improved rhythmicity in older (PR)_36_-expressing animals (Figure 6B). The absence of rescue could reflect cumulative toxicity from prolonged (PR)_36_ expression, a stronger deleterious effect of chronic TrpA1 activation, or effects attributable to age itself.

**Figure 6:**
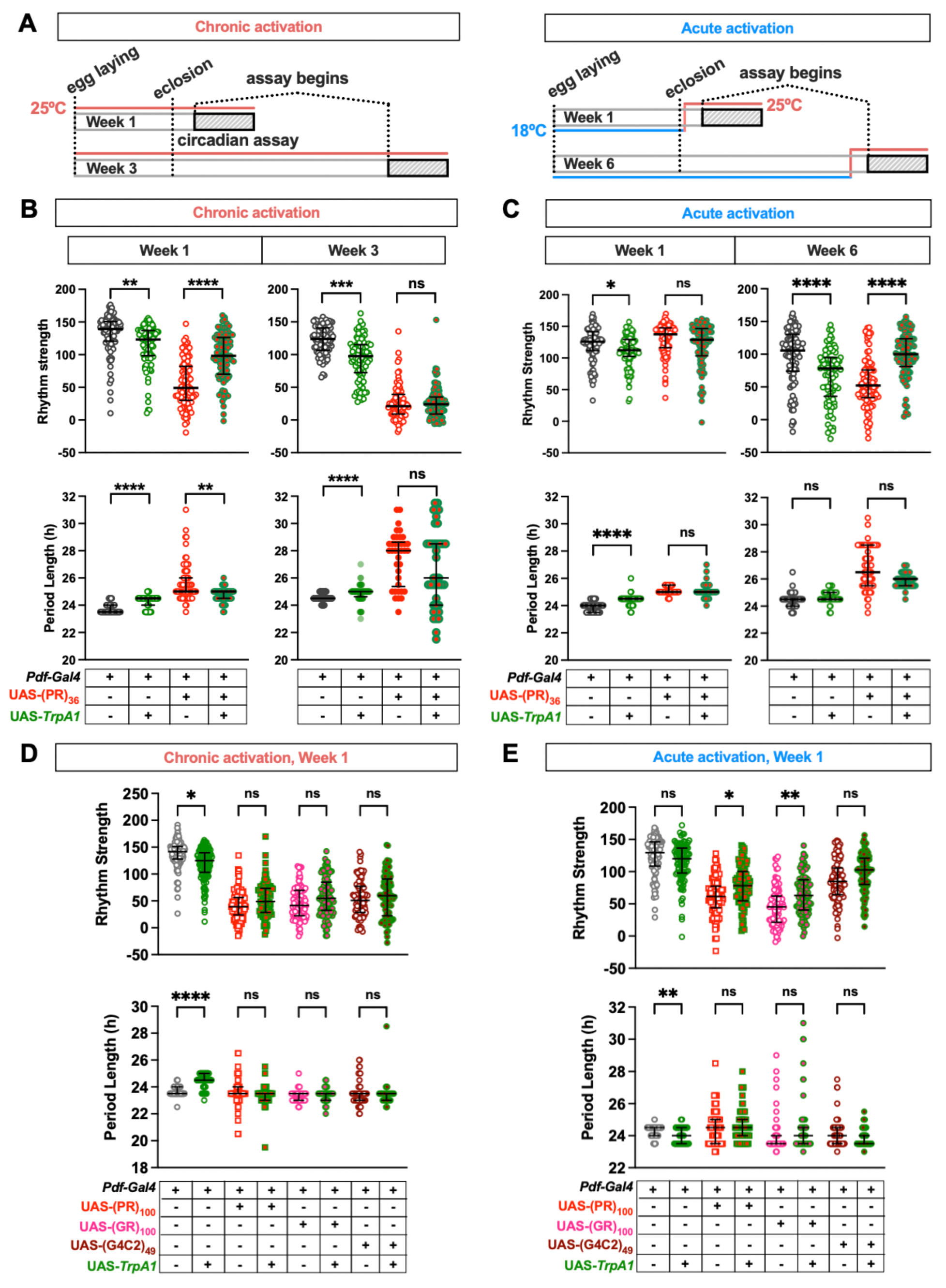
Increased neuronal activity can restore circadian function across ages and C9orf72 models. **(A)** Schematic of TrpA1 activation protocols. In the chronic activation condition (left), flies were raised and aged at 25°C from egg laying through the circadian assay. In the acute activation condition (right), flies were raised and aged at 18°C for either ∼1 day or ∼6 weeks, then shifted to 25°C for 3 days prior to the circadian assay to allow TrpA1 activation. Hatched black bars indicate the time window during which locomotor activity was recorded for circadian rhythm analysis. Bars and lines are not to scale. **(B-C)** Rhythm strength (top) and period length (bottom) in control males (*Pdf* > +) and males expressing TrpA1 alone, (PR)_36_ alone, or both. Results from chronic (A) and acute (B) TrpA1 activation are shown at 1, 3, and 6 weeks of age. Note that 3 weeks at 25°C and 6 weeks at 18°C correspond to approximately the same physiological age. The *Pdf* **> +** and *Pdf* > (PR)_36_ data from the chronic activation condition were previously shown in Figure 2 but were collected concurrently with the data from TrpA1-expressing genotypes and are included again to allow direct comparison. N = 92–96 (Chronic), 87-127 (Acute) for rhythm strength; n = 75–92 (Chronic, Week 1), 38–96 (Chronic, Week 3), 85–127 (Acute) for period length. **(D-E)** Rhythm strength (top) and period length (bottom) of control males (*Pdf > +*) and male flies expressing TrpA1 alone, indicated C9orf72 construct alone, or both. Results are shown for chronic (D) and acute (E) TrpA1 activation at 1 week of age. N = 87–126 (Chronic) and 94–96 (Acute) for rhythm strength; n = 63–125 (Chronic) and n = 63–96 (Acute) for period length. For all plots, data are presented as median and interquartile range. *p < 0.05, **p < 0.01, ***p < 0.001, ****p < 0.0001, ns: not significant; Kruskal-Wallis test followed by Dunn’s multiple comparisons test.

To test whether aging by itself impairs the capacity for neuronal activation to rescue (PR)_36_-induced dysfunction, we used an acute activation paradigm. Flies were reared and aged at 18°C, a temperature at which TrpA1 is inactive, and then shifted to 25°C 3 days prior to the start of the behavioral assay (Figure 6A). This allowed us to test the effects of adult-onset, acute TrpA1 activation. We conducted this experiment at two ages: approximately 1 week of age (flies that were 3–4 days old at the start of the circadian assay) and approximately 6 weeks of age (flies aged for 40 days at 18°C, then shifted to 25°C for 3 days before the assay). To estimate physiological age, we compared developmental timing at 18°C and 25°C and found that flies reached adulthood approximately 2.2 times faster at 25°C, consistent with previous reports^47,48^. Thus, 6-week-old flies reared at 18°C are roughly equivalent in physiological age to 3-week-old flies reared at 25°C. Our goal was not to precisely match physiological age, but rather to determine whether flies considerably older than 1 week of age can still benefit from increased neural activity.

Because the Gal4/UAS system drives lower transgene expression at 18°C compared to 25°C, (PR)_36_ had little behavioral effect in the 1-week-old group, preventing evaluation of TrpA1’s impact (Figure 6C). In contrast, by 6 weeks of age, (PR)_36_-expressing flies displayed a significant reduction in rhythm strength and longer period, comparable to 1-week olds at 25°C. Importantly, acute activation of TrpA1 at this stage rescued the (PR)_36_-induced rhythm defects, restoring rhythmicity and normalizing period (Figure 6C). These results indicate that aging per se does not prevent rescue of (PR)_36_-induced circadian dysfunction by increased neuronal excitability. Rather, the failure of TrpA1 co-expression to restore rhythmicity at 3 weeks under chronic activation conditions likely reflects cumulative toxicity of chronically elevated neuronal activation.

To test whether the rescue of C9orf72-associated circadian dysfunction by increased neuronal excitability depends on repeat type and length, we examined the effects of chronic TrpA1 activation on circadian deficits caused by (PR)_100_, (GR)_100_, and (G_4_C_2_)_49_ at 1 week of age. We found that continuous TrpA1 activation had little or no rescuing effect in these models (Figure 6D), possibly due to the greater toxicity of these constructs relative to (PR)_36_ (Figures 1 and 2). To assess whether reduced transgene expression would reveal a potential rescue window, we lowered C9orf72 construct expression by rearing animals at 18°C and applied the 1-week acute activation protocol described above. As expected, under these conditions the C9orf72 constructs caused milder circadian phenotypes than at 25°C (Figure 6E). Importantly, acute TrpA1 activation modestly but significantly improved rhythmicity in (PR)_100_- and (GR)_100_-expressing flies, and showed a slight, though not statistically significant, improvement in (G_4_C_2_)_49_ flies. These findings indicate that the ability of increased neuronal activity to rescue C9orf72-induced circadian dysfunction is not unique to (PR)_36_ but extends to other C9orf72 constructs when circadian deficits are relatively mild. However, the modest rescue observed, even under reduced expression levels and less severe phenotypes, suggests that the effectiveness of activity-based interventions depends on the specific conditions of activation. Together, these results show that enhancing neuronal activity can restore circadian function across multiple C9orf72 models and at different adult ages, provided that phenotypic severity is low, and activation is precisely tuned.

## Discussion

Disrupted circadian regulation is increasingly recognized as an early and potentially pathogenic feature of neurodegenerative disorders, including ALS and FTD^7,49^. Here, we used *Drosophila* to investigate how C9orf72-associated DPRs and HREs affect circadian behavior. Our data reveal that expression of (PR)_36_, (GR)_100_, or (G_4_C_2_)_49_ in PDF+ pacemaker neurons disrupted circadian rhythmicity and altered period length at ∼1 week of age, before detectable cell loss. At this early stage, the number of PDF+ s-LNvs remained unchanged, but PDF levels and the complexity of s-LNv dorsal projections were reduced, with severity correlating with behavioral phenotypes. Notably, the nature of circadian disruption varied by construct: (PR)_36_ lengthened period, while (GR)_100_ and (G_4_C_2_)_49_ shortened it, although the latter effect did not reach significance. Period lengthening may represent a less severe perturbation, while period shortening with reduced rhythmicity, may reflect more severe neuronal dysfunction. Supporting this idea, increasing PR repeat length and dosage, using (PR)_100_ and 2 copies of (PR)_36_, recapitulated the short-period phenotype observed with (GR)_100_ and (G_4_C_2_)_49_, while temporally restricted expression of (GR)_100_ or (G_4_C_2_)_49_ in adulthood induced long-period phenotypes similar to 1 copy of (PR)_36_. This variability in phenotypes provides a platform to investigate modifiers of circadian dysfunction at different stages of C9orf72 pathogenesis.

We also found that aging exacerbates circadian dysfunction. In both control and (PR)_36_-expressing flies, rhythm strength declined, and period length increased with age, with (PR)_36_ flies showing an earlier and more pronounced decline. By 6 weeks of age, some s-LNv loss was evident; however, the majority of C9orf72-expressing hemibrains retained all of these neurons, suggesting that neuronal dysfunction, rather than cell loss, underlies much of the observed behavioral decline. Interestingly, aging and increased PR dosage or longer repeats had opposing effects on period length; aging led to further lengthening, whereas higher dose and repeat length caused shortening. Prior work has shown that old flies lose behavioral rhythmicity despite retaining a robust molecular oscillator in central clock neurons^50^, suggesting that aging disproportionately disrupts circadian output pathways without affecting the central molecular clock. In contrast, C9orf72-associated toxicity may impair both output pathways and the central oscillator itself. Our findings underscore the complexity of interplay between genetic burden and aging in ALS/FTD-related circadian dysfunction.

Previous studies have reported both hyper- and hypoexcitability in C9orf72-ALS/FTD patient-derived neurons and mouse models^20–22,42^. In our *Drosophila* model, expression of (PR)_36_ reduced neural activity in PDF+ pacemaker neurons, as measured by the CaLexA calcium-dependent neural activity reporter. This reduction in neural activity likely contributes to the circadian behavioral deficits we observed. Supporting this interpretation, an increase in neuronal activity using the warmth-sensitive TrpA1 channel significantly rescued (PR)_36_-induced circadian dysfunction in young flies. When activated chronically at 25°C, a temperature that induces low-level TrpA1 activity, TrpA1 restored rhythm strength and normalized period length. The rescue effect of TrpA1 activation is unlikely to be mediated by restoration of PDF neuropeptide levels, as overexpression of PDF did not ameliorate C9orf72-induced deficits (Figure 4D). Further studies are needed to clarify the molecular mechanisms by which increased neuronal activity improves circadian function.

The efficacy of the activity-based rescue depended on phenotype severity and the parameters of TrpA1 activation. Chronic TrpA1 activation rescued (PR)_36_-induced deficits at 1 week but failed to do so at 3 weeks. Notably, TrpA1 activation alone reduced rhythmicity in older flies, suggesting that prolonged stimulation of PDF+ neurons becomes detrimental with age. To separate the effects of age from cumulative toxicity, we used an acute activation protocol in older flies reared at low temperature to suppress transgene expression until testing. Under these conditions, TrpA1 significantly improved circadian rhythmicity in (PR)_36_-expressing flies, indicating that age *per se* does not preclude rescue. Furthermore, acute neuronal activation at 1 week of age partially rescued circadian deficits caused by (PR)_100_ and (GR)_100_ expression, indicating that the restorative effect of increased neuronal activity extends beyond (PR)_36_-induced dysfunction. However, the magnitude of rescue was modest, underscoring that rescue efficacy is sensitive to the severity of underlying dysfunction, as well as to the timing, duration, and extent of neuronal activation.

The mixed findings of hyper- and hypoexcitability in mammalian C9orf72 models may be due to a biphasic trajectory, early hyperactivity followed by later hypoactivity as pathology advances. Several studies reported this transition in C9-ALS/FTD patient-derived induced pluripotent stem cell (iPSC) neurons^20,22,43^. These findings mirror clinical evidence that cortical and hippocampal hyperactivity, observed in early stages of Alzheimer’s disease, transitions to hypoactivity as neurodegeneration progresses^23–26^. In our *Drosophila* models, we found little evidence that increased neuronal activity exacerbates circadian deficits. Across multiple C9orf72 constructs and experimental conditions, TrpA1 activation either improved circadian rhythmicity or had little or no effect. Notably, in conditions where TrpA1 activation alone reduced rhythmicity in control flies, it did not further impair rhythmicity in C9orf72-expressing flies, even when there was room for additional decline. The effects of C9orf72 toxicity on neuronal excitability may be cell type–specific, producing sustained hypoexcitability in circadian pacemaker neurons across disease stages, while leading to a biphasic pattern in other neuronal populations. Our findings suggest that reduced excitability may contribute to dysfunction in some neural circuits, and that therapeutic benefit may depend on precisely calibrating excitability according to cell type, circuit function, and disease stage.

Our results offer broader relevance for neurodegenerative disease research. Circadian disturbances have been shown to precede motor or cognitive symptoms in Alzheimer’s and Parkinson’s diseases^51^, and disruptions in sleep-wake patterns have been observed in presymptomatic C9orf72 carriers and early-stage patients^7^. The delayed sleep onset in these patients is consistent with circadian period lengthening, although it may also reflect reduced sleep drive. Our results establish that C9orf72-linked toxicity can disrupt circadian function prior to cell loss, and that this disruption can be modulated through neural activity-based intervention, depending on several factors such as the severity of phenotypes and activation protocols. This highlights the value of circadian behavior as a sensitive readout of C9orf72 pathological processes and suggests the utility of *Drosophila* models for identifying novel cellular and genetic modifiers of C9orf72-linked pathology.

In conclusion, our study reveals a previously underappreciated vulnerability of the circadian system to C9orf72-associated toxicity. Expression of dipeptide and nucleotide repeats disrupts circadian behavior through a combination of reduced neuronal activity, impaired neuropeptide signaling, and structural alterations in pacemaker neuron projections, changes that emerge well before cell loss. Importantly, increasing neuronal activity can restore circadian function under select conditions. These findings link circadian dysfunction in C9orf72-ALS/FTD to decreased neuronal activity and suggest that treatments targeting neuronal excitability need to be matched to the type, stage, severity of dysfunction in individual patients.

### Limitations of the study

One limitation of this study is that the molecular mechanisms by which increased neuronal activity improves circadian function remain unclear. Future studies examining the molecular regulators of excitability and the downstream pathways modulated by neuronal activity may help identify the mechanisms underlying this rescue. Another limitation is that we focused on conditions with overt circadian phenotypes; thus, we cannot exclude the possibility that C9orf72-expressing circadian neurons pass through a transient hyperexcitable phase. Examination of additional models and activation protocols may be required to clarify whether an early phase of hyperexcitability contributes to disease progression. A third limitation is that temperature manipulations affected the duration of both TrpA1-mediated neuronal activation and C9orf72 expression in a correlated manner. Consequently, it is difficult to determine whether the key factor enabling increased neuronal activity to rescue C9orf72-associated circadian phenotypes in older flies is the shorter duration of activity itself or reduced C9orf72 expression resulting in a milder phenotype. Employing orthogonal expression systems with independent drivers for TrpA1 and C9orf72 would allow more precise dissection of their individual contributions.

## Methods

### Fly stocks

Flies were raised on a standard cornmeal-molasses-yeast medium and maintained under a 12-hour light:12-hour dark (LD) cycle at 25 °C unless otherwise noted. The following *Drosophila* lines were obtained from the Bloomington *Drosophila* Stock Center: *Pdf-Gal4* (#80939), UAS-(PR)_36_ (#58694), UAS-(GR)_100_ (#58696), UAS-(G_4_C_2_)_49_ (#84727), UAS-mCD8::GFP (#5137), UAS-*RedStinger* (#28281), UAS-*TrpA1* (#26263), UAS-*Kir2.1* (#91802), UAS-CaLexA (#66452) and the background control line *iso31* (*w*1118, #3605). *da-GS* line was generously provided by Dr. Amita Sehgal (University of Pennsylvania).

### Circadian behavior assays

For behavioral monitoring, adult flies were first entrained to a 12 h:12 h LD cycle at 25°C for at least three days. Flies were then individually housed in glass tubes containing 5% sucrose and 2% agar and monitored in constant darkness (DD) at 25°C for 6 days using the *Drosophila* Activity Monitoring (DAM) system (TriKinetics). Unless otherwise noted, flies were 3–4 days old at the start of the assay and are referred to as “1-week-olds.” Flies referred to as “3-week-olds” and “5-week-olds” were aged for 21–22 days and 35–36 days at 25°C, respectively, before the onset of DD monitoring. To induce the expression of (GR)_100_ and (G4C2)_49_ using the *da-GS* system, flies were fed 50 μM RU486 (mifepristone; Millipore Sigma). RU486 was diluted in 1% ethanol and added to a solution containing 5% sucrose and 2% agar, with 1% ethanol alone used as the vehicle control. For circadian experiments, flies were reared on standard *Drosophila* medium and then transferred to monitor tubes containing RU486 or vehicle solution. For acute TrpA1 activation, flies were raised and aged at 18°C to limit TrpA1 activity during development and aging. As expected with the Gal4/UAS system at lower temperatures, this condition also reduced (PR)_36_ expression. These flies were either 0–1 days old (“1-week-olds”) or 40–41 days old (“6-week-olds”) when transferred to 25°C for three days of activation prior to circadian assays at 25°C. Locomotor activity data were collected in 30-min bins and analyzed using Faas software (developed by M. Boudinot and F. Rouyer, Institut des Neurosciences Paris-Saclay). Rhythm strength was calculated for all flies, while period length was determined only for those classified as rhythmic, defined by a rhythm strength threshold of ≥25, as described in more detail previously^52^.

### Immunohistochemistry

Following dissection, adult brains were fixed in 1% paraformaldehyde for 45 min at room temperature. The samples were blocked in 1% normal goat serum (NGS) for PDF, GFP, and RFP antibody staining. The primary and secondary antibodies were diluted in 5% NGS and incubated overnight at 4°C. The primary antibodies utilized were mouse monoclonal anti-PDF antibody (Developmental Studies Hybridoma Bank, University of Iowa, Cat# PDF C7) at a dilution of 1:500, and rabbit GFP polyclonal antibody (Invitrogen, Thermo Fisher Scientific, Cat# A-11122) and rat monoclonal anti-RFP antibody (ChromoTek, Cat# 5f8) at a dilution of 1:1000. The secondary antibodies used were goat anti-mouse IgG (H+L) cross-adsorbed secondary antibody, Cyanine5 (Invitrogen, Thermo Fisher Scientific, Cat# A10524), goat anti-rabbit IgG (H+L) cross-adsorbed secondary antibody, Alexa Fluor 488 (Invitrogen, Thermo Fisher Scientific, Cat# A11008) and goat anti-rat IgG (H+L) cross-adsorbed secondary antibody, Alexa Fluor 647 (Invitrogen, Thermo Fisher Scientific, Cat# A-21247) all at 1:1000. RedStinger signal in dissected brains was visualized without antibody staining after fixation and washes. All images were captured using a Leica SP8 confocal microscope with 1 μm Z-steps. Identical acquisition settings were used across genotypes within each experiment.

### Image quantification and statistical analysis

Image analysis was performed in Fiji/ImageJ (https://fiji.sc) by experimenters blinded to genotype. For each marker (PDF, CD8::GFP, RFP, and CaLexA), we measured total fluorescence intensity from maximum intensity projections and subtracted background signal to quantify expression levels. To assess projection complexity, Scholl analysis was performed on maximum intensity projections of the s-LNv dorsal arbors. Concentric circles spaced 10 µm apart were centered at the anatomical bend where the dorsal projections make an approximately 90-degree turn, and the number of intersections between the projections and each circle was quantified, as described ^40^. All statistical analyses were performed using Prism 10 (GraphPad Software). Group comparisons were made using the Kruskal-Wallis test with Dunn’s T3 correction for multiple comparisons. Data distributions were assessed with the Kolmogorov-Smirnov and Shapiro-Wilk tests. Given that some datasets deviated from normality, non-parametric methods were uniformly applied across all analyses for consistency.

## Resource availability

### Lead contact

Further information and requests for resources and reagents should be directed to and will be fulfilled by the lead contact, Kyunghee Koh.

### Materials availability

This study did not generate new materials.

### Data and code availability

Data: Any additional information required to reanalyze the data reported in this article is available from the lead contact upon request.

Code: This study does not report original code.

## Acknowledgments

We thank the Bloomington *Drosophila* Stock Center and Dr. Amita Sehgal for providing fly stocks and M. Boudinot and Francis Rouyer for the Faas software. We are grateful to Drs. Pierra Pasinelli, Hristelina Ilieva, and Brigid Jensen, members of the Jefferson Weinberg ALS Center, and Liz Matthews for their valuable feedback. This work was supported by a postdoctoral fellowship award from Japanese Society for Promotion of Science (G24KJ0033 and G25K18771 to S.I.), grants from the NIH/NINDS (R21NS135762 to K.K and RF1NS114128-01A1S1 to A.R.H., D.T., and K.K.), and by funding from Thomas Jefferson University through the Jefferson Center for Synaptic Biology Awards (to K.K.).

## Author contributions

Conceptualization: S.I., B.P.J., O.A., D.T., A.R.H., and K.K; methodology: S.I., B.P.J., O.A., and K.K; investigation: S.I., B.P.J., O.A., S.G., and A.S.; writing – original draft, S.I., O.A., and K.K.; writing – review and editing, all authors; supervision: A.K. and K.K.; project administration: K.K.; and funding acquisition, S.I., D.T., A.R.H., and K.K.

## Declaration of generative AI and AI-assisted technologies

During the preparation of this work, the authors used ChatGPT to improve clarity and flow without altering the content. The authors take full responsibility for the content of the publication.

